# Evolution of *Enterococcus faecium* to A Combination of Daptomycin and Fosfomycin Reveals Distinct and Diverse Adaptive Strategies

**DOI:** 10.1101/2021.12.20.473606

**Authors:** Adeline Supandy, Heer H. Mehta, Truc T. Tran, William R. Miller, Rutan Zhang, Libin Xu, Cesar A. Arias, Yousif Shamoo

**Affiliations:** Department of Biosciences, Rice University, Houston, TX, USA; Center for Infectious Diseases Research, Houston Methodist Research Institute, Houston, TX, USA; Division of Infectious Diseases, Houston Methodist Hospital, Houston, TX, USA; Department of Medicinal Chemistry, University of Washington, Seattle, WA, USA

## Abstract

Infections caused by vancomycin-resistant *Enterococcus faecium* (VREfm) are an important public health threat. VREfm have become increasingly resistant to the front-line antibiotic, daptomycin (DAP). As such, the use of DAP combination therapies (like fosfomycin [FOS]), has received increased attention. Antibiotic combinations could extend the efficacy of current available antibiotics and potentially delay the onset of further resistance. We investigated the potential for *E. faecium* HOU503, a clinical VREfm isolate that is DAP and FOS susceptible, to develop resistance to a DAP-FOS combination. Of particular interest was whether the genetic drivers for DAP-FOS resistance might be epistatic and, thus, potentially decrease the efficacy of a combinatorial approach in either inhibiting VREfm or in delaying the onset of resistance. We show that resistance to DAP-FOS could be achieved by independent mutations to proteins responsible for cell wall synthesis for FOS and in altering membrane dynamics for DAP. However, we did not observe genetic drivers that exhibited substantial cross-drug epistasis that could undermine DAP-FOS combination. Of interest was that FOS resistance in HOU503 was largely mediated by changes in phosphoenolpyruvate (PEP) flux as a result of mutations in pyruvate kinase (*pyk*). Increasing PEP flux could be a readily accessible mechanism for FOS resistance in many pathogens. Importantly, we show that HOU503 were able to develop DAP resistance through a variety of biochemical mechanisms and were able to employ different adaptive strategies. Finally, we showed that the addition of FOS can prolong the efficacy of DAP, significantly extending the timeline to resistance *in vitro*.

**Importance:** While the discovery of antibiotics was one of the greatest health care advances in history, its success is being challenged by the emergence of multidrug-resistant pathogens, including vancomycin-resistant enterococci (VRE). Daptomycin (DAP), a lipopeptide antibiotic that targets cell membrane, is currently prescribed as a frontline drug to treat VRE infections. However, emergence of daptomycin-resistant VRE is concerning. Consequently, DAP-Fosfomycin (FOS) combination (DF) has been proposed as a potential method to maintain DAP efficacy. Here, we provide evidence that DF successfully delayed the emergence of resistance *in vitro*. Genetic data indicates that resistance was acquired independently, with little evidence of significant cross-drug epistasis that could undermine a combinatorial approach. We also uncovered a novel FOS resistance mechanism, through changes in phosphoenolpyruvate (PEP) flux, that may potentially be shared with other bacterial species. Additionally, we also have evidence showing that *E. faecium* was able to employ different resistance mechanisms.

## Introduction

The enormous success of antibiotics is being challenged by the rise of increasingly resistant pathogens that threaten to return us to a pre-antibiotic era with potentially devasting consequences (1). The complexity of the antibiotic resistance problem is perhaps best manifested by the increasing number of multi-drug resistant (MDR) pathogens that include vancomycin-resistant enterococci (VRE) and, increasingly, vancomycin and daptomycin-resistant enterococci (VDRE) (2, 3). Frequently, physicians prescribe daptomycin (DAP) for serious VRE infections (2, 4, 5). DAP acts by inserting itself into the cell membrane of Gram-positive bacteria in a calcium-dependent manner, causing the displacement of critical cell membrane proteins and cell death (6, 7). DAP insertion into the cell membrane is mediated by binding to the anionic phospholipids phosphatidylglycerol (PG) as well as lipid II precursors (7). To date, the two main strategies employed by enterococci in DAP resistance include: 1) an increase in cell envelope charge to decrease the amount of DAP binding (repulsion), which is commonly seen in *S. aureus* (8–10) and 2) the rearrangement of membrane architecture (redistribution), which is commonly seen in *E. faecalis* (11). Genetic changes associated with the redistribution phenotype in *E. faecalis* occurred in the the LiaFSR system, a stress response system involved with cell envelope homeostasis, and cardiolipin synthases (Cls), a protein involved in the synthesis of cardiolipin which is a constituent of the bacterial cell membrane (12–14).

Interestingly, *E. faecium* exhibit a more diverse ensemble of DAP resistance mechanisms compared to *E. faecalis*. While mutations in the LiaFSR pathway remain commonly seen, mutations in YycFG, a two-component system pathway involved in cell wall metabolism and restructuring, have also been observed (12–17). Additionally, a previous study also identified mutations in the *yvcRS* operon, which caused an increase in cell surface charge and decreased DAP binding (13).

Due to the emergence of DAP resistance, new approaches to restore clinical efficacy of DAP have been proposed (2, 13, 14, 18, 19). Combining DAP with other antibiotics is an attractive approach (20–25). Fosfomycin (FOS) has recently joined the list of potential partner with DAP as DAP-FOS combination (DF) and have already shown promising results against enterococcal infections (26–30). FOS is an antibiotic that acts as an analog of phosphoenolpyruvate (PEP) to inhibit the activity of MurAA, which is involved in the first committed step of peptidoglycan synthesis (31). As a drug, FOS has been approved for clinical use orally in the US and has been used to treat *E. faecium* infections. A study by a group in China has discovered that clinical FOS-R *E. faecium* isolates possess the *fosB3* gene, a variant of an enzyme that inactivates FOS through chemical modifications. (32, 33). Another study also showed that clinical VREfm isolates acquire FOS resistance due to a cysteine to aspartate mutation in the active site of MurAA (34). These studies show that *E. faecium* can develop FOS resistance through both mutational resistance and acquisition of exogenous resistance determinants.

In this study, we use *in vitro* experimental evolution to identify evolutionary trajectories that lead to DF resistance. Using the clinical strain *E. faecium* HOU503, we showed that resistance to the DF combination was achieved via independent mutations conferring resistance to each drug separately. Moreover, while DF did not prove synergistic against HOU503, this combination did significantly extend the timeline by which *E. faecium* became resistant to DAP. This delay suggests that DF combinatorial therapy could extend the efficacy of both drugs and delay the rise of resistant variants.

## Materials and Methods

### Bacterial strain and growth conditions

*E. faecium* HOU503 is a VRE clinical strain that harbors common LiaRS substitutions (LiaR^W73C^ and LiaS^T120A^) with DAP and FOS susceptibility (DAP MIC = 3 μg/ml, FOS MIC = 128 μg/ml) (35, 36). FOS MIC breakpoint was extrapolated from the CLSI breakpoint for *E. faecalis*. All cultures were grown in Brain Heart Infusion (BHI) at 37°C and shaking at 225 rpm unless noted otherwise. 50 mg/L CaCl_2_ was used unless noted otherwise. Glucose-6-Phosphate was not added as it did not create noticeable differences in the FOS MIC values (Fig. S1 in supplemental material).

### *In vitro* adaptation of *E. faecium* HOU503 to DAP and FOS

*E. faecium* HOU503 was adapted to DAP and FOS individually as well as in combination with five populations per condition. The evolution was conducted in the following manner. Single colonies were used to start the adaptation. Each day, a 1% dilution of the culture with best growth was passaged to a new tube with increasing antibiotic concentration. Serial passage was continued until resistant colonies (MIC ≥ 2x CLSI breakpoint) were obtained. One population served as a no-drug control. At the end of adaptation, each population was serially diluted onto non-selective BHI plates. Three isolates from each population were selected at random for further analysis. The evolution was conducted in duplicate. A difference between the two replicate is the antibiotic selection gradient, where the first replicate (FT1) experienced slower increase in the antibiotic concentration compared to the second replicate (FT2).

### Determination of DAP to FOS ratio using a checkerboard assay

A 96-well plate containing the first antibiotic (DAP) along the ordinate and the second antibiotic (FOS) along the abscissa was prepared. Inoculum was standardized to OD_600_ of 0.05. The plates were incubated overnight. FIC Index value was calculated according to Chen et al., 2014 (32).

### Determination of end-point-isolates fitness and MIC

Fitness of each end-point-isolate was tested by comparing the growth curve of each resistant isolate to HOU503. Briefly, overnight cultures were normalized to an OD_600_ of 0.05 and used to inoculate fresh BHI in a microplate. OD_600_ was measured every 5 min at 37°C for 24 h in BioTek EpochII microplate reader. The DAP and FOS MICs of all end-point-isolates were tested using agar dilution assay in BHI agar with CaCl_2_ (37). Both experiments were conducted in biological triplicate.

### Determination of PEP concentration

PEP quantification was performed with the PEP Colorimetric/Fluorometric Assay Kit (Sigma-Aldrich) according to the manufacturer’s instructions with modifications. Briefly, mid-log phase cultures were pelleted and lysed using the bead mill method. Resulting supernatant was used for both PEP and Bradford assay. Subsequent steps were done as directed by the manufacturer protocol. The assay was conducted in half-area 96-well plates and fluorescence was measured in BioTek EpochII microplate reader at 570 nm. The data were normalized to protein concentration obtained from Bradford assay. Assay was conducted in biological triplicate and technical duplicate.

### Determination of mRNA expression by qPCR

Total RNA was extracted using the Qiagen RNeasy Mini kit with additional incubation of sample with 5 μL of 5 U/mL mutanolysin and 12.5 μL of 200 mg/mL lysozyme at 37°C for 30 min. Samples were then treated with TURBO™ DNase according to manufacturer’s protocol and the ThermoFisher SuperScript III kit was used to synthesize cDNA. qPCR was performed according to manufacturer’s protocol using the BioRad iQ™ SYBR Green Master Mix on a Bio-Rad CFX connect real-time system. *gyrB* was used as the internal reference. Relative expression was calculated using the 2^−ΔΔCT^ method. This assay was conducted in biological and technical triplicate. Primer sequences are shown in Table S1.

### Membrane phospholipid distribution and BODIPY-DAP binding

The membrane dye 10 *N*-nonyl acridine orange (NAO) is used to visualize anionic phospholipids according to a previously published protocol with several modifications (11). Briefly, cells were grown to mid-log phase in TSB and then incubated with 250 nM NAO at 37°C with shaking in the dark for 3.5 h. Cells were then washed in saline three times and immobilized on 1% agarose pad.

Conjugation of Bodipy-FL (BDP) to DAP was carried out as described previously (38). Briefly, BDP and DAP were incubated in 0.2 M sodium carbonate buffer, pH 8.5, with shaking at room temperature for 1 h. This was followed by dialysis against distilled H_2_O at 4°C overnight. The resulting BDP:DAP was used as a proxy to visualize binding of DAP to cells according to previously published protocol (11, 13, 39). Cells were grown until mid-log phase in BHI with CaCl_2_. Then, cells were incubated with BDP:DAP in the dark and shaking at 37°C for 20 min. After staining, cells were washed once with HEPES (20 mM, pH 7.0) and immobilized on 1% agarose pad.

Fluorescence images of both assays were taking on Keyence BZ-Z710 microscope with 100x objective using the FITC filter (excitation 490 nm, emission 528 nm). Both assays were conducted in triplicate.

### Membrane fluidity analysis by *in vivo* Laurdan generalized polarization spectroscopy

Laurdan, 6-dodecanoyl-2-dimethylaminonaphthalene, is a membrane dye used to determine alterations of membrane fluidity (40). Laurdan generalized polarization (GP) spectroscopy was performed according to a previously published protocol (41). Briefly, mid-log phase culture in BHI supplemented with 0.1% glucose was incubated with 10 μM Laurdan in a light-protected box with shaking at 37°C for 10 min. The sample was then washed four times with phosphate buffered saline (PBS) supplemented with 0.1% glucose and CaCl_2_. After the final wash, the sample was resuspended in PBS/glucose/calcium solution. Assay was conducted with four technical replicates of each condition (untreated and ethanol) and two replicates of control Laurdan background. Baseline fluorescence intensity was measured with excitation at 350 nm and emission at 435 nm and 500 nm for 5 min at 37°C on a Synergy H1 spectrophotometer (BioTek). Cells were then treated with 10% ethanol as a membrane fluidizer. Measurements taken every 2 minutes as above for the next 15 min. The Laurdan GP index was calculated with the formula (*I*_435_ – *I*_500_)/(*I*_435_ + *I*_500_), where *I* is the fluorescent intensity at the specified wavelength.

### Lipid Extraction for lipidomics

Lipid extraction was carried out as described previously with minor modifications (42). Briefly, cells were grown without the presence of antibiotics before they were pelleted and processed. Additionally, dried extracts were reconstituted in 500 μL of 1:1 chloroform-methanol solution mix. For analysis, 10 μl of the lipid extract per 10 mg of original pellets was transferred to an LC vial, dried under Argon or Nitrogen, and reconstituted to 100 μL with 2:1 acetonitrile-methanol solution mix.

### Membrane lipid compositions by hydrophilic-interaction liquid chromatography (HILIC) Ion Mobility-Mass Spectrometry (IM-MS)

For liquid chromatography, bacterial lipids were separated by a Waters UPLC (Waters Corp., Milford, MA, USA) as described previously (42–44).

The Waters Synapt XS platform was used for lipidomics analysis with similar parameters as described in Zhang et al., 2021 (42)

### Data Analysis for HILIC-IM-IS data

Data alignment, chromatographic peaks detection, and normalization were performed in Progenesis QI (Nonlinear Dynamics). A pooled quality control sample was used as the alignment reference. The default “All Compounds” method of normalization was used to correct for variation in the total ion current amongst samples. Lipid identifications were made based on m/z (within 10 ppm mass accuracy), retention time, and CCS with an in-house version of LipidPioneer, modified to contain the major lipid species, including free fatty acids (FFAs), DGDGs, PGs, CLs, and LysylPGs with fatty acyl compositions ranging from 25:0 to 38:0 (total carbons:total degree unsaturation), and LiPydomics (44–46). The quantitation of FFAs was performed based on their ion mobility retained chromatographic peaks using Waters DriftScope 2.5 and TargetLynx (Waters Corp., Milford, MA, USA).

### Whole genome sequencing (WGS) and analysis

Genomic DNA of HOU503 and end-point-isolates were isolated using the Qiagen DNeasy UltraClean Microbial Kit following manufacturer’s instructions with additional incubation at 37°C in the microbead solution with 5 μL of 5 U/mL mutanolysin and 12.5 μL of 200 mg/mL lysozyme for 1 h. Paired-end libraries were generated using the Plexwell™ 384 library generation kit. Samples were sequenced by a commercial facility with 2×150 bp reads on Hi-seq. End-point-isolates were sequenced with a minimum coverage of 100x while populations were sequenced with at least 300x coverage. Illumina sequences were compared to the closed ancestor genomes (503F_del_LiaR under bioproject PRJNA544687) using the Breseq genomic pipeline. Population samples were processed using the polymorphism (-p) mode. All genomic sequences are under PRJNA789547.

## Results

### Adaptation to DAP-FOS combination successfully delays the onset of resistance

To study the effect of a DF combination against *E. faecium*, we evolved *E. faecium* HOU503 to either each drug individually or in combination (Fig. 1).

**FIG 1.**
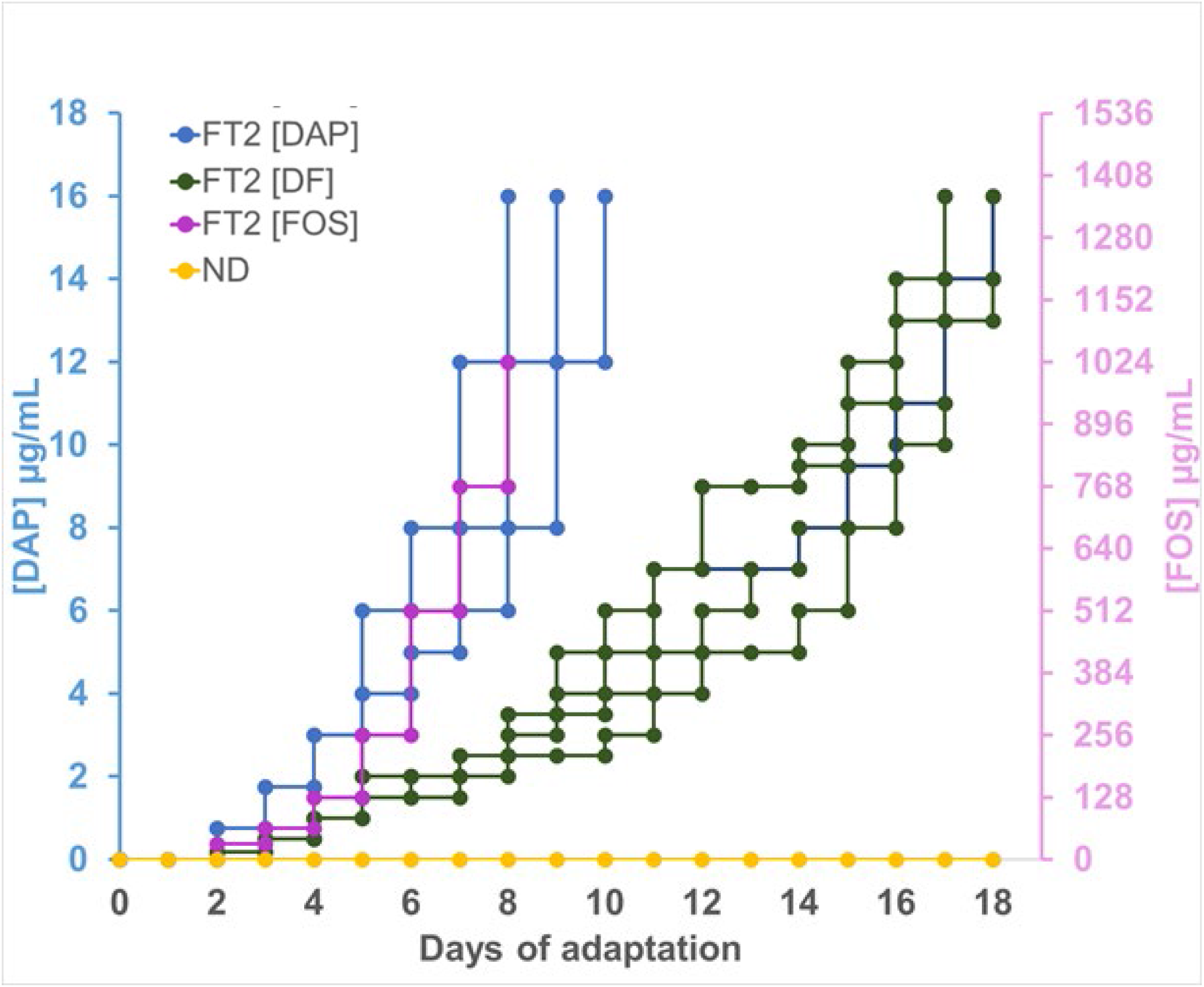
Adaptation rate of HOU503 to DAP, FOS, and DF of FT2. 5 populations of HOU503 were adapted daily to increasing either DAP, FOS, or DF concentrations until resistance was achieved. Blue, purple, and green lines denote DAP, FOS, and DF adaptation, respectively. Rate of HOU503 exposure to DAP, FOS, and DF in FT1 can be found in Fig. S3 in the supplemental material.

The DAP to FOS ratio was determined using a checkerboard assay. Since the DF combination did not exhibit synergistic activity when tested against HOU503 (Fig. S2 in supplemental material), the DAP:FOS ratio was set to 1:85.3 μg/mL according to the checkerboard assay result. Ten HOU503 populations were evolved to DAP, FOS, and DF in two groups of five (FT1 and FT2) using *in vitro* experimental evolution (13, 47–49). FT1 and FT2 used two distinct selection gradients. At the end of adaptation, end-point-isolates were selected randomly for whole-genome-sequencing (WGS) to identify mutations acquired and further characterization (Table 1-3).

**Table 1.** Whole genome sequencing data of DAP-evolved HOU503 isolates^*a*^

|  |  | DAP MIC | FOS MIC | *pgsA* | *cls* | *gdpD* | CDP-AP | *murF* | pyruvate carboxylase | Additional Mutations |
| --- | --- | --- | --- | --- | --- | --- | --- | --- | --- | --- |
| FT1 | D1 | 64 | 128 |  | H215R;<br>R211L |  |  |  |  | +4 |
|  | D2 | 64 | 128 |  | H215R;<br>R211L |  |  |  |  | +4 |
|  | D3 | 64 | 128 |  | H215R;<br>R211L |  |  |  |  | +4 |
|  | D4 | 64 | 128 |  |  |  | -62 | S323R |  | +6 |
|  | D5 | 64 | 128 |  |  |  | -62 | S323R |  | +4 |
|  | D6 | 64 | 170.7 |  | A20D |  |  |  |  | +4 |
|  | D7 | 26.7 | 128 | A98D |  |  |  |  |  | +3 |
|  | D8 | 32 | 128 |  |  |  |  |  |  | +5 |
|  | D9 | 16 | 128 |  | A20D |  |  |  |  | +2 |
|  | D10 | 64 | 128 |  |  |  | -45 |  | Q1073* | +7 |
|  | D11 | 32 | 128 |  | A20D |  |  |  | Q1073* | +6 |
|  | D12 | 9.3 | 128 | V85F |  |  |  |  | Q1073* | +3 |
|  | D13 | 32 | 170.7 |  | R211L |  |  |  |  | +4 |
|  | D14 | 13.3 | 128 |  |  |  | -62 |  |  | +4 |
|  | D15 | 32 | 213.3 |  | H215R |  |  |  |  | +3 |
| FT2 | D16 | 32 | 128 | T70P |  |  |  |  |  | +2 |
|  | D17 | 26.7 | 128 | T70P |  |  |  |  |  | +3 |
|  | D18 | 13.3 | 64 | T70P |  |  |  |  |  | +2 |
|  | D19 | 42.7 | 64 |  | R211Q |  |  |  |  | +3 |
|  | D20 | 32 | 128 |  | R211Q |  |  |  |  | +3 |
|  | D21 | 32 | 64 |  | R211Q |  |  |  |  | +3 |
|  | D22 | >128 | 128 |  |  | L93H | -62 |  |  | +3 |
|  | D23 | >128 | 128 |  |  | L93H | -62 |  |  | +3 |
|  | D24 | 128 | 128 |  |  | L93H | -62 |  |  | +4 |
|  | D25 | 21.3 | 128 |  | H215R |  |  |  |  | +2 |
|  | D26 | 16 | 64 |  | H215R |  |  |  |  | +2 |
|  | D27 | 16 | 64 |  | H215R |  |  |  |  | +2 |
|  | D28 | 64 | 128 |  |  |  | -62 |  |  | +3 |
|  | D29 | 64 | 128 |  |  |  | -62 |  |  | +3 |
|  | D30 | 64 | 64 |  |  |  | -62 |  |  | +3 |

<sup>a</sup>The selected mutations shown do not include those observed in the no-drug control isolates or initial starting ancestor. MIC reported are averaged over three biological replicates. The “Additional Mutations” column include occurrence(s) of *van* operon (*vanRSHAXYZ*) loss. D5 and D6 retained one *van* operon copy while other isolates lost both copies. HOU503 (LiaR<sup>W73C</sup> and LiaS<sup>T120A</sup>) DAP MIC: 3 µg/mL; FOS MIC: 128 µg/mL.

**Table 2.** Whole genome sequencing data of FOS-evolved HOU503 isolates^*b*^

|  |  | DAP MIC | FOS MIC | *pyk* | MurAB | Additional Mutations |
| --- | --- | --- | --- | --- | --- | --- |
| FT1 | F1 | 5.3 | >1024 | G135S | -73 | +4 |
|  | F2 | 4 | >1024 | K165R | -73 | +3 |
|  | F3 | 8 | >3584 | P330L | -73 | +2 |
|  | F4 | 4 | >1024 | P12S | -73 | +4 |
|  | F5 | 5.3 | >1024 | G162R | -73 | +3 |
|  | F6 | 4 | >1024 | P330L | -73 | +5 |
|  | F7 | 6.7 | >1024 | P266A | Insertion upstream | +2 |
|  | F8 | 5.3 | >1024 | P266A | Insertion upstream | +2 |
|  | F9 | 4 | >1024 | P266A | Insertion upstream | +2 |
|  | F10 | 6.7 | >1024 | M289I | -73 | +4 |
|  | F11 | 6.7 | >1024 | M289I | -73 | +4 |
|  | F12 | 5.3 | >1024 | M289I | -73 | +4 |
|  | F13 | 12 | 512 |  | -73 | +4 |
|  | F14 | 10.7 | 597.3 |  | -73 | +3 |
|  | F15 | 5.3 | 768 |  | -73 | +3 |
| FT2 | F16 | 4 | >3584 | L264P | -73 | +2 |
|  | F17 | 4 | 512 |  | -73 | +3 |
|  | F18 | 4 | >1024 | G257S | -73 | +5 |
|  | F19 | 4 | >1024 | M319I | -73 | +4 |
|  | F20 | 13.3 | >1024 | G235R | -73 | +3 |
|  | F21 | 13.3 | >1024 | G235R | -73 | +4 |
|  | F22 | 2.7 | >1024 | M256I | -73 | +3 |
|  | F23 | 4 | >1024 | R298L | -73 | +2 |
|  | F24 | 8 | >1024 | R298L | -73 | +3 |
|  | F25 | 8 | >1024 | R298L | -73 | +2 |
|  | F26 | 4 | >1024 | M289I | -73 | +5 |
|  | F27 | 4 | >1024 | E303G | -73 | +2 |
|  | F28 | 5.3 | >1024 | P297R | -73 | +3 |

<sup>b</sup>The selected mutations shown do not include those observed in the no-drug control isolates or initial starting ancestor. Two isolates were not included due to contamination issues. The “Additional Mutations” column include occurrence(s) of *van* operon (*vanRSHAXYZ*) loss. All isolates lost both *van* operon copies. MIC reported are averaged over three biological replicates. HOU503 (LiaR<sup>W73C</sup> and LiaS<sup>T120A</sup>) DAP MIC: 3 µg/mL; FOS MIC: 128 µg/mL.

**Table 3.**
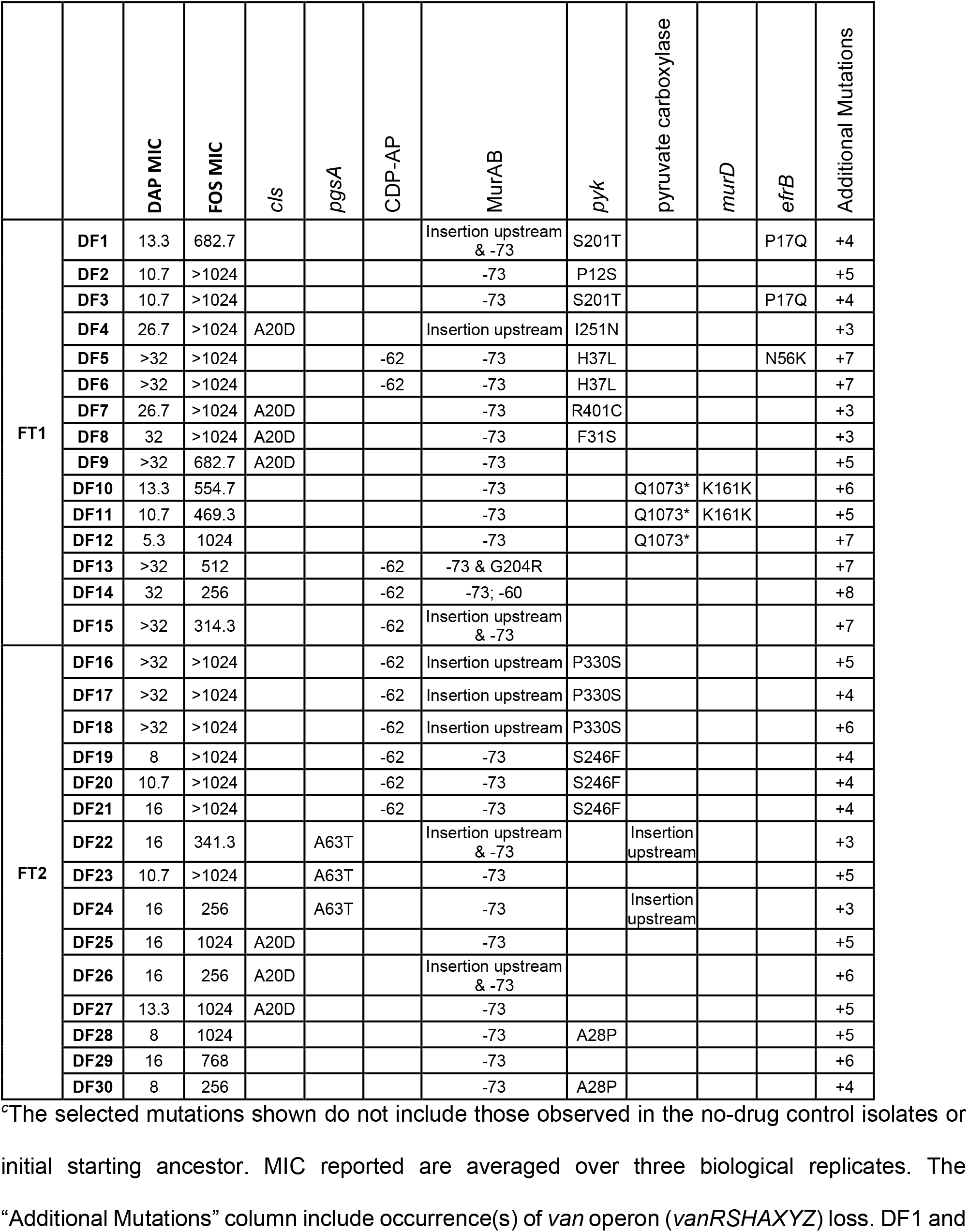
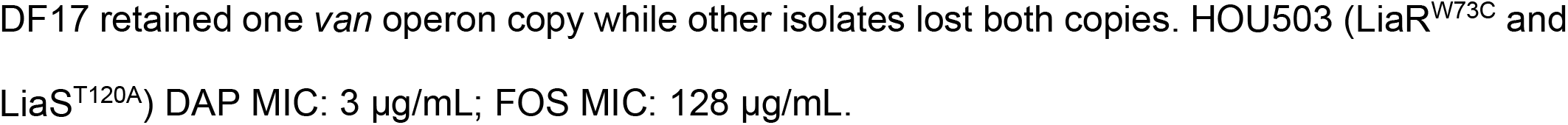
Whole genome sequencing data of DF-evolved HOU503 isolates^*c*^

Importantly, HOU503 experienced more difficulty in adapting to DF combination (FT2 shown in Fig. 1). The DF combination resulted in a longer timeline to DF resistance, about 50% longer compared to the populations adapted to each drug individually. If the antibiotic concentration increments were lower then, as expected, DAP and FOS resistance within the populations increased more readily, but the DF combination still delayed DAP resistance (Fig. S3 in supplemental material).

### Increased MurAB expression as an important step in FOS resistance

FOS has generally been used against Gram-negative bacteria, but it has returned to prominence due to its potential as a “companion” drug against a wide range of Gram-positive MDR pathogens, including enterococci (27, 50).

In contrast to previous studies, all HOU503 FOS-R isolates evolved mutation(s) affecting MurAB, an isozyme of MurAA, either as a single nucleotide polymorphism (SNP) and/or an insertion of a transposon upstream of the gene (Table 2 and 3) (28, 32). In *S. aureus*, under FOS attack, *murAB* has been shown to be upregulated and complement MurAA in maintaining cell wall synthesis (51). To verify if a similar mechanism was employed by HOU503, RT-qPCR was performed. Due to the challenges associated with engineering mutations in *E. faecium* to create single mutants, FOS resistant isolates from FOS evolved populations, F5 (MurAB^-73^*pyk*^G126R^) and F8 (MurAB^Isrt^*pyk*^P266A^) were used. Indeed, regardless of the type of mutation, *murAB* expression was significantly upregulated compared to the ancestor (Fig. 2A). The ubiquitous presence of mutations upstream of MurAB suggests that MurAB may play a critical role in FOS resistance in *E. faecium* (Table 2 and 3).

**FIG 2.**
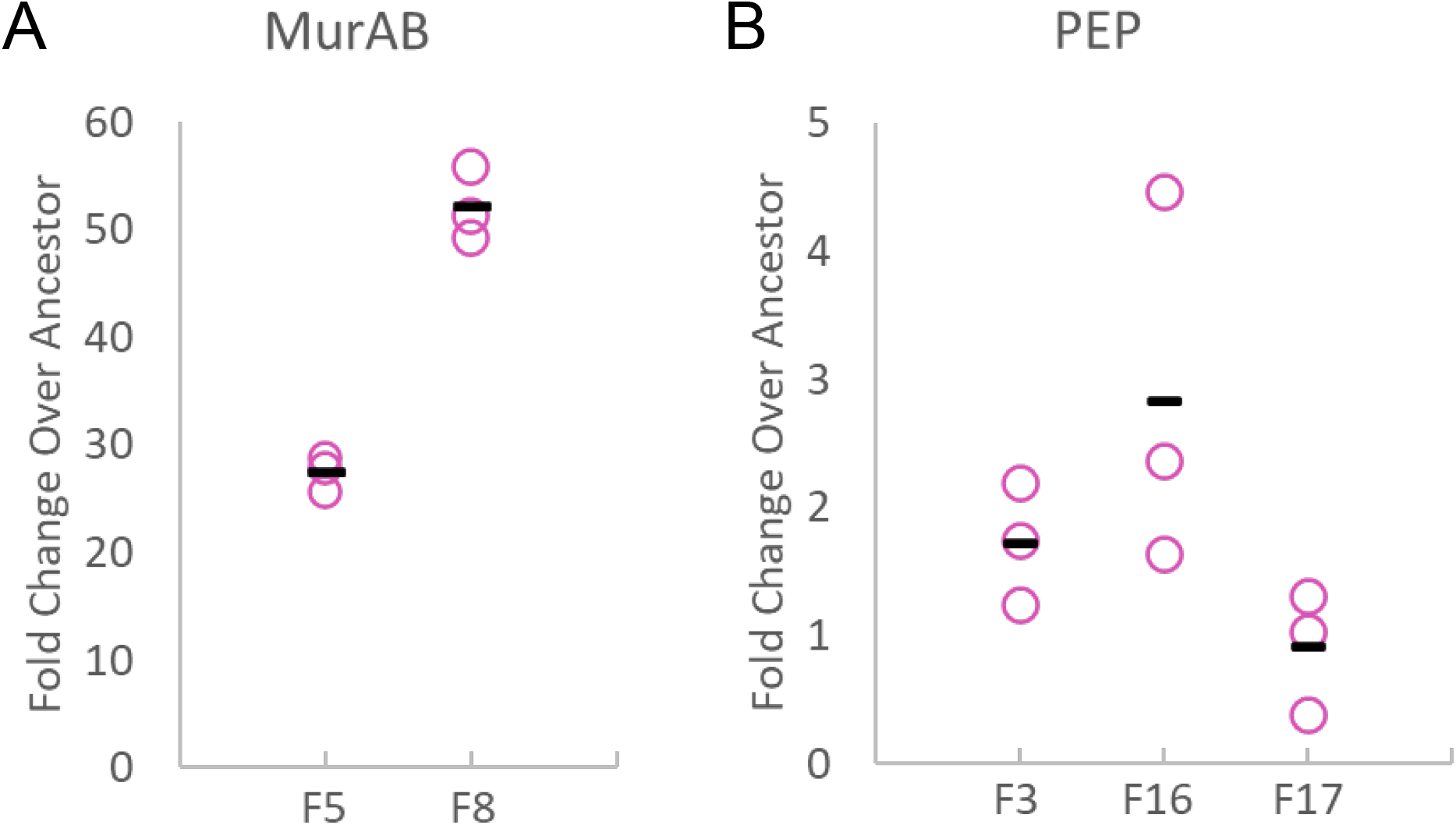
FOS resistance mediated by upregulation of MurAB expression and PEP. (A) qPCR was performed to measure changes in MurAB transcript expression. Assay was conducted in biological triplicates. Housekeeping gene: *gyrB* (B) Phosphoenol pyruvate (PEP) concentration was measured through the PEP Colorimetric/Fluorometric Assay Kit (Sigma-Aldrich). Experiment was conducted in biological triplicates with technical duplicates.

### Increased PEP level as a mechanism of FOS resistance

The other mutation facilitating FOS resistance is a mutation in pyruvate kinase (*pyk*), an enzyme involved in production of pyruvate from PEP (Table 2 and 3). As PEP is the substrate for MurAA, and a structural analog of FOS, we postulated that the mutations in *pyk* decrease the flux of PEP to pyruvate, allowing more free PEP to compete against FOS for the MurAA active site. To address this hypothesis, an assay measuring PEP level was conducted on selected isolates with and without *pyk* mutations.

Isolates that include a *pyk* mutation, F3 (MurAB^-73^*pyk*^P330L^) and F16 (MurAB^-73^*pyk*^L264P^), showed about a 2-fold increase of PEP concentration. Conversely, F17 (MurAB^-73^), an isolate without *pyk* mutations, has similar levels of PEP as the ancestor (Fig. 2B). This result suggests that adaptive mutations in *pyk* probably decreases its activity resulting in a net increase in free PEP, leading to increased FOS resistance.

Moreover, there is a significant difference in FOS MIC between isolates with and without mutations in *pyk*. The presence of *pyk* mutations increased the FOS MIC of isolates by more than 7-fold (from 512 to >3584 μg/mL) (Table 2). Interestingly, the effect of PEP in FOS resistance seems to be concentration dependent as the isolate with more PEP, F16, exhibited a shorter lag time during growth studies than F3 during FOS stress (Fig. S4 in supplemental material). Overall, these results suggest that adaptive mutations in *pyk* can act as important drivers for *E. faecium* to achieve higher FOS resistance.

### HOU503 achieved DAP resistance through biochemically distinct adaptive strategies

Previous studies on DAP resistance in enterococci have identified adaptive changes consistent with membrane composition remodeling (11, 52, 53). Indeed, with background mutations in LiaRS, DAP-R HOU503 isolates gained mutations in genes involved in membrane lipid synthesis (CDP-AP, *pgsA, cls*, and *gdpD*) (Table 1). Taken together, these mutations suggest that DAP resistance in HOU503 was facilitated by changes to cell membrane lipid flux. However, changes in the homeostasis of a single lipid species can have pleiotropic effects on other lipid pools and membrane fluidity, which have recently been implicated in the antimicrobial action of DAP (6, 10, 44). To provide a more holistic assessment of lipid homeostasis during adaptation to DAP we quantitated changes in: 1) membrane lipid pools (lipidomic analyses), 2) membrane fluidity (Laurdan dye assay); 3) anionic lipid displacement (NAO staining); and 4) DAP binding (BDP:DAP staining).

As expected, PG levels play an important role in DAP resistance. D19 that includes *cls*^R211Q^ showed a significant increase in the amount of cardiolipin, but lower PG levels compared to the ancestor (Fig. 3A and B). This is consistent with a previous study that showed DAP adaptive mutations within *cls* cause increased cardiolipin synthesis activity (52). Additionally, as cardiolipin is synthesized from PG, its increased activity would also explain the significantly lower amount of PG (Fig. 3A). These changes in PG and CL levels extended across the range of lipid species consistent with generalized changes in the PG/CL pools. HOU503 also seems to attain increased DAP resistance through decreased PgsA activity as seen in D16. As PgsA catalyzes the formation of PG precursor, its decreased activity could explain the lower PG amount in D16 (*pgsA*^T70P^) (Fig. 3A). Intriguingly, strains harboring either *pgsA*^T70P^ or *cls*^R11Q^ correlate with lipid patterning and localization consistent with a redistribution mechanism in which anionic phospholipids are moved away from the division septa (Fig. 4B).

**FIG 3.**
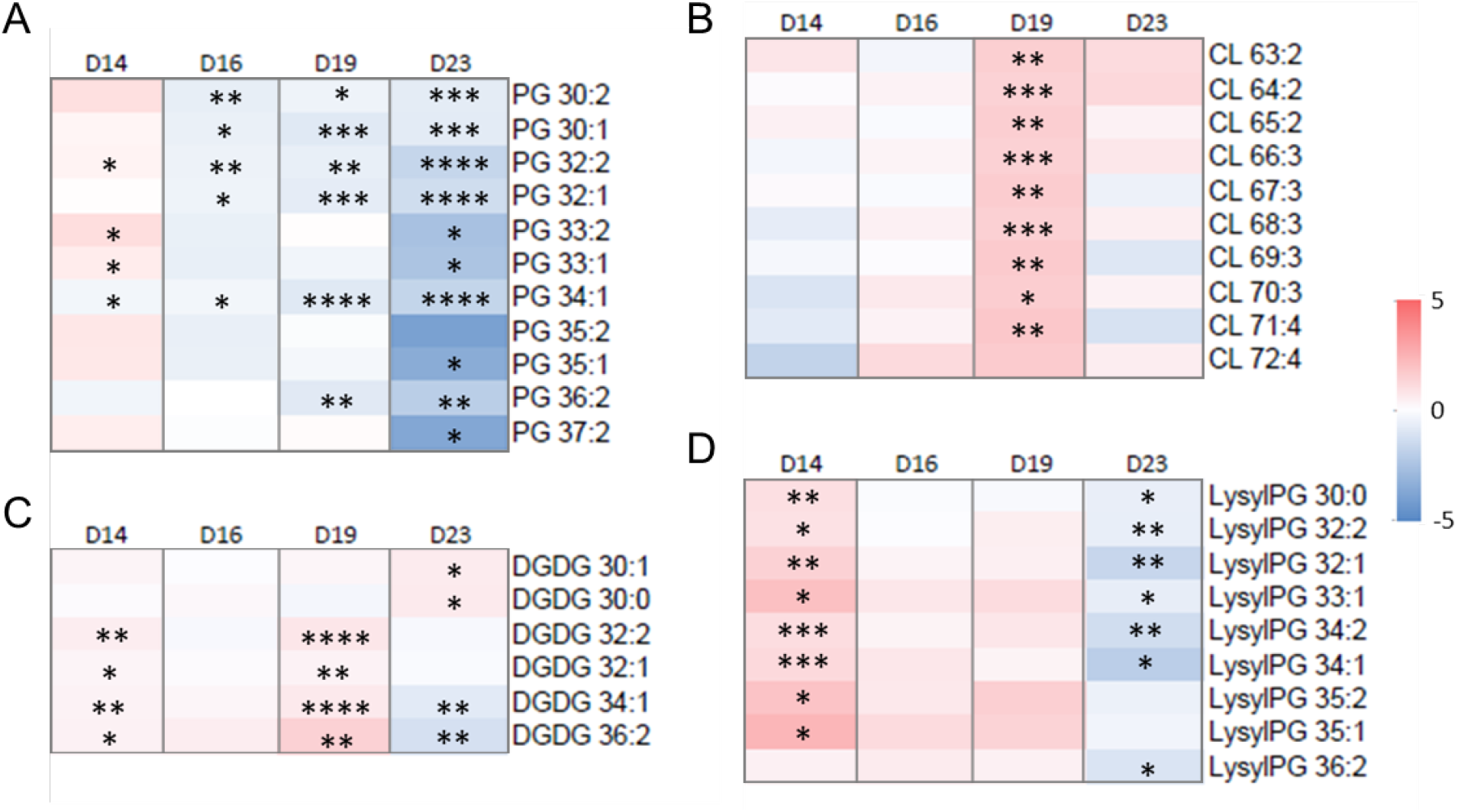
DAP resistance in HOU503 is correlated with changes to membrane lipid composition. Abundance of (A) phosphatidylglycerol (PGs), (B) cardiolipins (CLs), (C) diglucosyl-diacylglycerols (DGDGs) and (D) lysl-phosphatidylglycerol (LysylPGs) were normalized to dry bacterial pellet weight and internal standard ion abundance. Individual lipid species are represented as the number of carbons: the degree of unsaturation in the fatty acid chains. Figure presented as heatmap of log_2_ fold change over ancestor. Significance was calculated with Student’s t-test. *: P ≤ 0.05; **: P ≤ 0.01; ***: P ≤ 0.001; ****: P ≤ 0.0001. For primary data, refer to Fig. S5 in the supplementary material.

**FIG 4.**
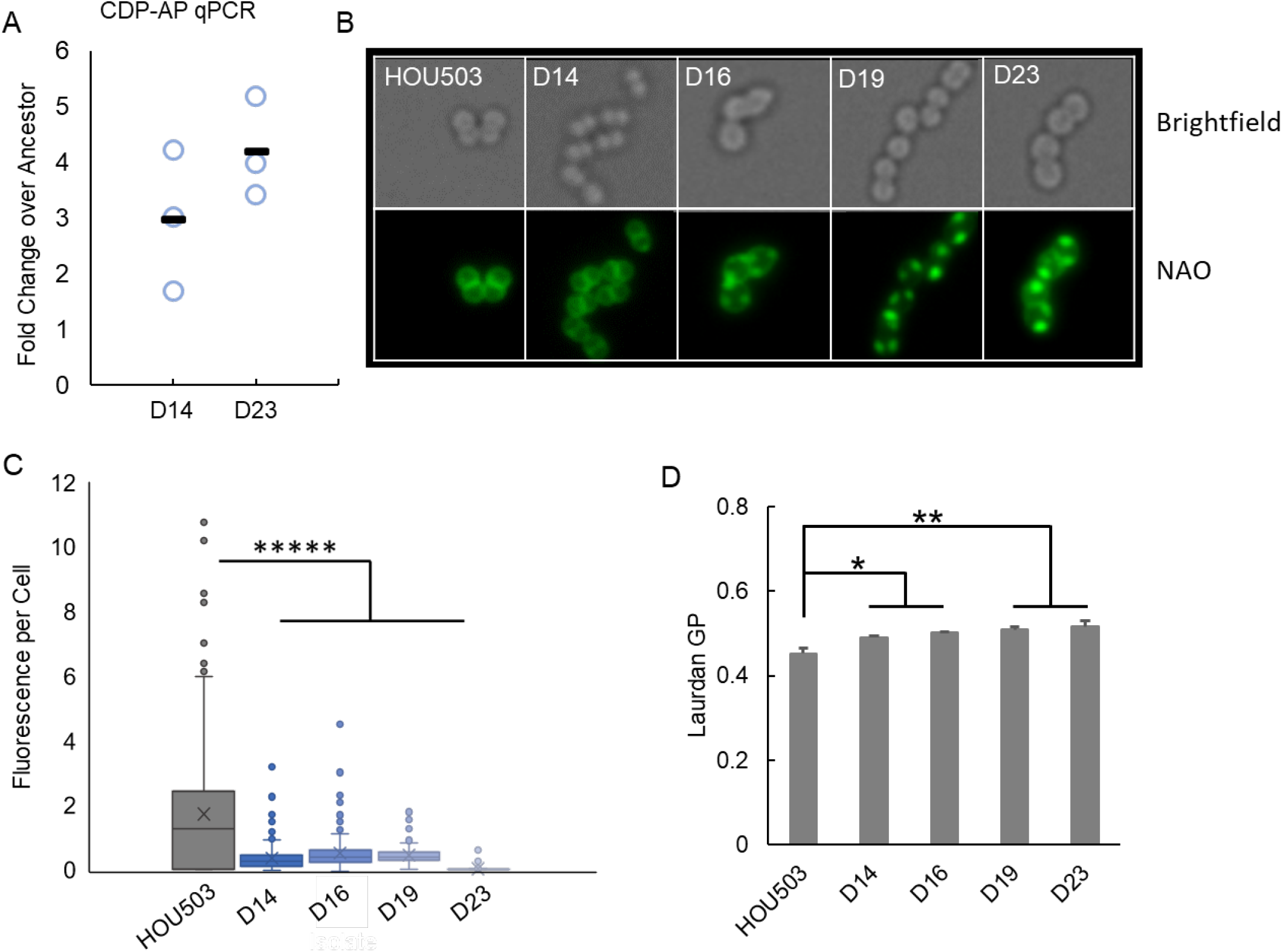
DAP resistance in *E. faecium* HOU503 displays diverse phenotypes consistent with the identified biochemical mechanisms. (A) Increased CDP-AP transcript expression is observed in mutants with a SNP located upstream of CDP-AP. (B) Visualization of anionic phospholipids in the cell membrane using NAO (250 nM) staining of representative cells of HOU503 and its DAP-R isolates. Top images in the panel show brightfield while bottom row show fluorescent images. (C) Quantification of Bodipy-labelled DAP (BDP:DAP) binding of HOU503 and selected DAP-R derivatives when stained with 8 μg/mL BDP:DAP. (See also Fig. S6 in the supplemental material.) (D) Quantification of the mean baseline fluidity as calculated through the fluorescent Laurdan dye assay for HOU503 and its DAP-R derivatives. (See also Fig. S7 in the supplemental material.) Significance for (C) and (D) were calculated using the Mann-Whitney test with post hoc Holm Bonferroni adjustment and Student’s t-test, respectively.. *: P ≤ 0.05; **: P ≤ 0.01; ***: P ≤ 0.001; ****: P ≤ 0.0001.

Mutations were also identified upstream of a gene annotated as a CDP-AP family protein (FGF98_RS11550), but whose function is unknown (Table 1). CDP-AP family proteins include proteins involved in the biosynthesis of essential cell membrane lipids (54). As the mutation occurred upstream of the putative CDP-AP gene, we carried out RT-qPCR and found a 3 to 5-fold increase in expression among isolates carrying the mutation (D14 and D23) (Fig. 4A). D14 (CDP-AP^-62^) has a significantly greater amount of most species of PG and lysyl-PG compared to the ancestor (Fig. 3A and 3D, respectively). NAO staining of D14 does not suggest phospholipid redistribution but is more consistent with electrostatic repulsion phenotype in leading to increased DAP resistance (Fig. 4B).

Interestingly, when CDP-AP^-62^ is paired with adaptive changes in *gdpD*, DAP resistance markedly increases. D23 (CDP-AP^-62^*gdpD*^L93H^) has a significantly lower amount of both PG and lysyl-PG (Fig. 3A and 3D, respectively) and exhibited a more redistribution-like phenotype with a rearrangement in the cell membrane architecture (Fig. 4B). Moreover, the addition of *gdpD*^L93H^ also boosted the DAP MIC by >9-fold (13.3 vs >128 μg/mL) (Table 1).

Significantly, despite the variety of mutations these isolates carry, all tested isolates have a significantly more rigid membrane and bound significantly less BDP:DAP than the ancestor HOU503 (Fig. 4C-D). This decrease in membrane fluidity may influence the way and amount of DAP that can insert itself into the membrane thus leading to decreased BDP:DAP binding and overall greater resistance to DAP “attack”. In general, these data showed that although *E. faecium* HOU503 can utilize multiple pathways in becoming DAP resistant, an overall changes in PG and more rigid membrane seem to be consistent phenotypes (6, 7).

### HOU503 evolved resistance to DF combination through independent pathways

As mentioned above, DAP and FOS affect the cell membrane and cell wall, respectively. One of the reasons we chose to study DF combination was to determine if the genetic drivers for DF resistance might be epistatic and thus potentially decrease the efficacy of a combinatorial approach in either inhibiting VREfm or delaying the onset of DAP resistance. As shown in Table 3, we did not observe any evidence of substantial cross-drug epistasis that could undermine a combinatorial approach. Changes within populations adapting to the single drug separately did not markedly increase their resistance to the other. Instead, we saw that resistance to DF was achieved independently with mutations in cell wall synthesis and central metabolism for FOS and in altering cell membrane lipid dynamics for DAP (Fig. 5).

**FIG 5.**
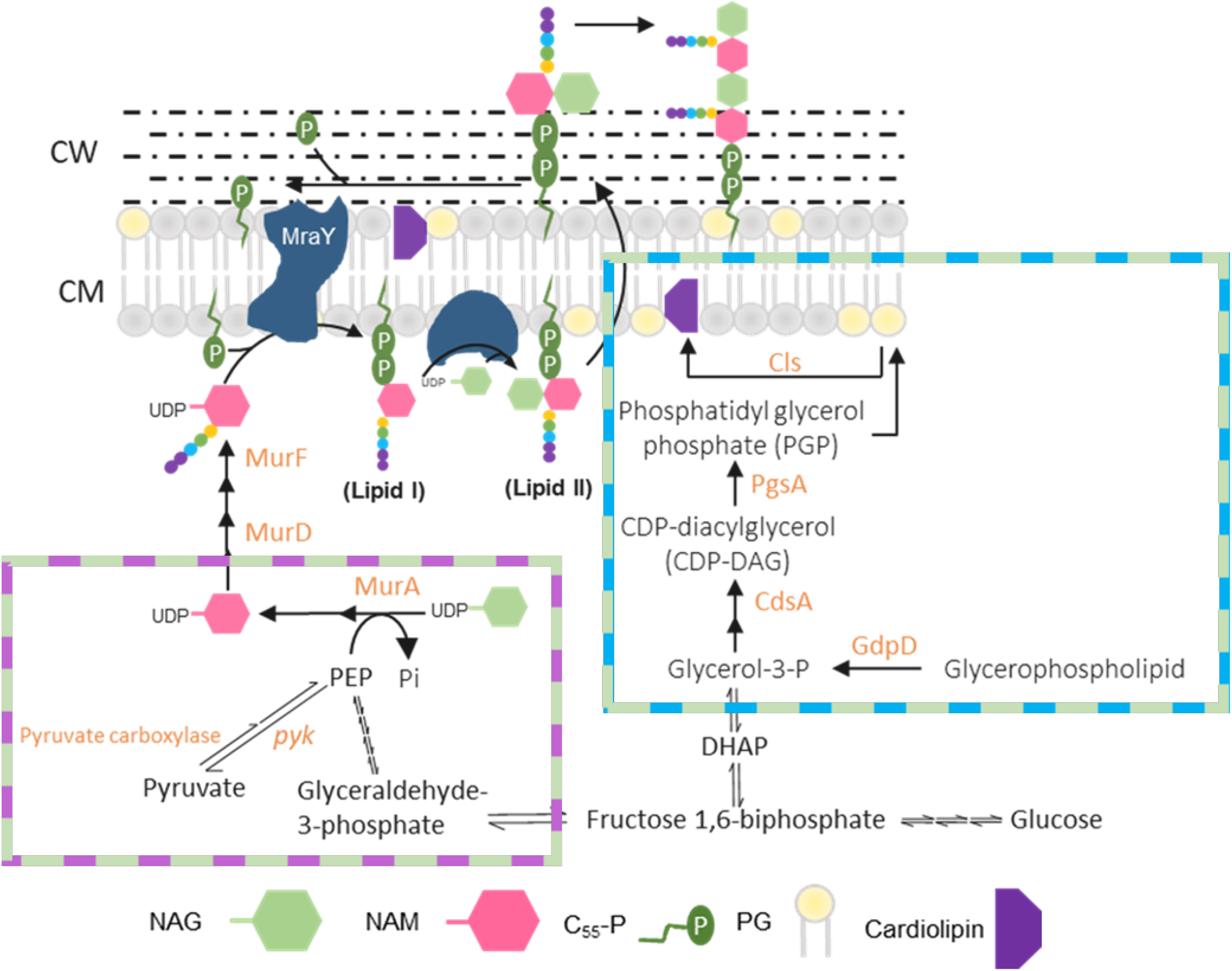
Schematic of changes in the enterococcal cell envelope synthesis pathways associated with DAP, FOS, and DF resistance. Cell wall synthesis begins with the synthesis of the uridine diphosphate-N-acetylmuramic acid (UDP-NAM) pentatpeptide peptidoglycan precursor by the sequential action of MurA to MurF. UDP-NAM pentapeptide is then transferred to the lipid carrier C_55_P forming lipid I, which is then used to form lipid II. Meanwhile, the cell membrane lipids phosphatidylglycerol (PG) and cardiolipin are synthesized through numerous precursors and various proteins, including those in the CDP-AP family. These two pathways are joined through the central metabolism pathway. DAP resistance was mainly mediated by mutations to proteins involved in the membrane lipid synthesis pathway (blue box) while FOS resistance was facilitated through mutations in the cell wall synthesis and central metabolism pathway (purple box). Meanwhile, DF resistance was achieved independently through changes to both pathways separately (green boxes). Adapted from Hancock et al., 2014 and Grein et al., 2021. Created with Biorender.

## Discussion

MDR pathogens like VRE are increasingly responsible for difficult to treat infections that require the use of drugs-of-last resort such as DAP. Due to the occurrence of DAP-resistant *E. faecium* infections, strategies to prolong or augment the activity of this antibiotic, such as combination therapy, are needed (5, 55). FOS has been shown to exhibit synergy with DAP against other Gram-positive bacterial infections (50, 56). We investigated a DAP and FOS drug combination against a clinical strain of *E. faecium* to discern whether the combination may delay the onset of DAP resistance or, in the worst-case scenario, facilitate the quick ascent of strains with resistance to both.

Although the DF combination did not exhibit synergistic activity against HOU503, the onset of DAP resistance was successfully delayed. Our *in vitro* evolution studies showed that DF resistance in HOU503 occurred independently with no clear evidence of mutations that could act as epistatic genetic drivers providing resistance to both drugs simultaneously. The lack of cross-drug epistasis also supports the potential efficacy of DF combinatorial approach in either treating VREfm infections and delaying the onset of DAP resistance.

Previous studies have shown that DAP resistance in enterococci can be achieved through “repulsion” of the antibiotic molecule or the rearrangement of the cell membrane architecture (8, 9, 11). Interestingly, *E. faecium* has shown its capability to utilize either mechanism, which is influenced by several factors, including genetic background and adaptive environment (13). In this paper, we showed that DAP-resistant *E. faecium* isolates derived from the same ancestor and evolved in the same environment were able to utilize both resistance strategies.

Isolates that exhibited a “repulsion”-like mechanism acquired a mutation upstream of a CDP-AP family protein (FGF98_RS11550), which include those involved in the biosynthesis of cell membrane lipids (54). While the net effect of changes in the regulation of this CDP-AP produce a phenotype similar to that of *mprF*, which catalyzes the addition of lysyl group to PG and has been previously implicated in the repulsion phenotype, CDP-AP shares less than 8% homology to either *mprF1* or *mprF2* suggesting a distinct biochemical role (10, 13). Moreover, D14 also showed no evidence of phospholipid redistribution. Interestingly, an extra mutation in *gdpD* in D23 (CDP-AP^-62^*gdpD*^L93H^) reverses the phenotype of the resistance mechanism to be more redistribution-like with a rearrangement in the cell membrane architecture. Moreover, when paired with mutation to *gdpD*, the DAP MIC increases by at least 9-fold to over 128 μg/ml.

Furthermore, HOU503 also exhibited cell membrane architecture rearrangement when it acquired mutations in either *cls* or *pgsA*. The observed *cls* mutation is consistent with our earlier study suggesting that adaptive changes in *cls* in response to DAP are correlated with an increase in Cls activity (52). Although insufficient by itself, adaptive mutations in *cls* when combined with pre-existing mutations in the LiaFSR regulon seem to be sufficient for DAP resistance (57). On the other hand, mutations in *pgsA* seem to decrease its enzymatic activity, leading to a decrease in PG content. This decrease in available PG likely may contribute to resistance by providing less opportunity for the critical PG-dependent DAP binding event that is part of the initial DAP interaction with the cell membrane (7).

Remarkably, despite the different adaptative mechanisms seen in DAP-resistant *E. faecium*, each of the end-point-isolates bound less BDP:DAP compared to the ancestor. Additionally, although the DAP-R isolates have different membrane lipid compositions, they all show evidence for a more rigid cell membrane than the ancestor. Both strains that included upregulation of CDP-AP (D14 and D23) showed larger changes in lipids with longer chain lengths and may be a clue into the potential role or substrate preference of CDP-AP (Fig. 3). As membrane fluidity has been implicated previously in DAP binding, it is possible that these changes in membrane rigidity decrease DAP binding to the cell leading to less BDP:DAP being bound (6).

In contrast, mutations observed during adaptation to FOS were less diverse. FOS acts by competing with PEP for the MurAA active site and the mutations observed were focused on either MurAB, a homolog of MurAA, or pyruvate kinase (*pyk)*. The mutations observed in *pyk* are likely decrease-of-function mutations as intrinsic PEP levels in isolates with *pyk* mutations were higher than the ancestor. Conversely, mutations upstream of MurAB resulted in a significant upregulation of MurAB expression. A study in *S. aureus* showed that MurAB was able to compensate for, or complement, MurAA function (51). Our qPCR data suggests that HOU503 may have employed a similar mechanism. However, this hypothesis requires further studies to be validated, especially regarding whether MurAB can catalyze the same reaction as MurAA. Mutations upstream of MurAB were ubiquitous across all our populations suggesting that it is a critical contributor to FOS resistance. Interestingly, while HOU503 possess the *fosB3* gene (FGF98_01880), which was implicated in *E. faecium* FOS resistance previously, we did not observe any mutations affecting this gene (28, 32). Additionally, we also did not observe mutations affecting MurAA which had been documented previously in FOS-R VREfm isolates (34). The lack of mutation within MurAA in DF-R isolates was surprising. MurAA is a promising target to achieve FOS resistance (34). Additionally, mutations in MurAA has been seen previously in DAP-R *E. faecium* isolate lacking the *liaR* response regulator, suggesting it may contribute to cellular fitness during DAP exposure (47). This absence implies that resistance strategies heavily rely on the genetic background of the strain.

In summary, we show that *E. faecium* evolved resistance to DF combination independently, without evidence of epistatic mutations that affect both drugs simultaneously (Fig. 5). DAP resistance in HOU503 was mediated by changes to the cell membrane lipid profiles and fluidity. Despite the different mutations observed, overall changes to PG content and higher membrane rigidity were shared among all the DAP-resistant isolates tested. This suggest that these may be the consistent phenotypes for DAP-resistant *E. faecium* HOU503. On the other hand, the increase in MurAB expression and PEP levels were fundamental to FOS resistance in HOU503. Despite lacking synergistic activity, DF combination was able to extend the timeline in HOU503 to achieve resistance quite significantly, signifying its potential for the treatment of *E. faecium* infections clinically.

## Acknowledgments

This work was supported by National Institutes of Health, National Institute of Allergy and Infectious Diseases grants R01A1080714 to Y.S., K08 AI135093-01A1 to W.R.M., R01AI136979 to L.X., R01-AI148342, R01-AI134637, P01-AI152999, and K24-AI121296 to C.A.A. Funding agencies did not play a role in experimental design, performance, or analysis. C.A.A. has received grants from Merck, MeMEd Diagnostics, and Entasis Therapeutics. W.R.M. has received a grant from Merck, and honoraria from Achaogen and Shionogi. T.T. has received a grant from Merck.

